# Temporally resolved single cell transcriptomics in a human model of amniogenesis

**DOI:** 10.1101/2023.09.07.556553

**Authors:** Nikola Sekulovski, Amber E. Carleton, Anusha A. Rengarajan, Chien-Wei Lin, Lauren N. Juga, Allison E. Whorton, Jenna K. Schmidt, Thaddeus G. Golos, Kenichiro Taniguchi

**Author notes:** Corresponding author: **CONTACT:** Kenichiro Taniguchi, 8701 Watertown Plank Road, B4245 BSB CB062, Milwaukee, WI 53226. These authors contributed equally to this work (Nikola Sekulovski, Amber E. Carleton, Anusha A. Rengarajan, Chien-Wei Lin).

## Abstract

Amniogenesis is triggered in a collection of pluripotent epiblast cells as the human embryo implants. To gain insights into the critical but poorly understood transcriptional machinery governing amnion fate determination, we examined the evolving transcriptome of a developing human pluripotent stem cell-derived amnion model at the single cell level. This analysis revealed several continuous amniotic fate progressing states with state-specific markers, which include a previously unrecognized CLDN10^+^ amnion progenitor state. Strikingly, we found that expression of CLDN10 is restricted to the amnion-epiblast boundary region in the human post-implantation amniotic sac model as well as in a peri-gastrula cynomolgus macaque embryo, bolstering the growing notion that, at this stage, the amnion-epiblast boundary is a site of active amniogenesis. Bioinformatic analysis of published primate peri-gastrula single cell sequencing data further confirmed that CLDN10 is expressed in cells progressing to amnion. Additionally, our loss of function analysis shows that CLDN10 promotes amniotic but suppresses primordial germ cell-like fate. Overall, this study presents a comprehensive amniogenic single cell transcriptomic resource and identifies a previously unrecognized CLDN10^+^ amnion progenitor population at the amnion-epiblast boundary of the primate peri-gastrula.

## INTRODUCTION

Amniogenesis is initiated during implantation in humans, and leads to the formation of an amniotic sac structure that surrounds and protects the developing embryo (Enders et al., 1983; Enders et al., 1986; Miki et al., 2005; Sasaki et al., 2016; Shahbazi and Zernicka-Goetz, 2018; Taniguchi et al., 2019). At implantation, the embryo (referred to at this time as a blastocyst) contains three morphologically and molecularly distinct cell types: 1) a collection of unpolarized pluripotent epiblast cells, precursors to both the embryo proper and the amniotic ectoderm, 2) a surrounding layer of polarized trophectoderm, a placental tissue precursor, and 3) an underlying extraembryonic primitive endoderm, a yolk sac precursor. Upon implantation, the pluripotent epiblast cells initiate epithelial polarization to form a cyst with a central lumen, the future amniotic cavity (Carleton et al., 2022; Shao and Fu, 2022). This event is followed by the fate transition of pluripotent epiblast cells that are in close proximity to the uterus to squamous amniotic ectoderm, forming a clear boundary between amnion and pluripotent epiblast territories of the cyst (Shahbazi and Zernicka-Goetz, 2018; Shao and Fu, 2022; Taniguchi et al., 2019). This asymmetric amnion-epiblast structure, known as the amniotic sac, provides the foundation for the next essential developmental steps (e.g., primitive streak formation, neural specification).

Recent studies using early human and monkey embryos have provided a basic understanding of the transcriptomic characteristics of early primate amniogenesis (Bergmann et al., 2022; Nakamura et al., 2016; Nakamura et al., 2017; Sasaki et al., 2016; Tyser et al., 2021; Yang et al., 2021). Yet, experimental dissection of the molecular mechanisms involved in this process is difficult in these *in vivo* models. To enable molecular and cellular investigations, we and others developed human pluripotent stem cell (hPSC)-derived amnion models and showed that BMP signaling plays a crucial role in initiating amniogenesis (Chen et al., 2021; Overeem et al., 2023; Shao et al., 2017a; Shao et al., 2017b; Zheng et al., 2019). Recently, we further explored the BMP-dependent amniogenic transcriptional cascade, identifying several distinct transcriptional stages, as well as a requirement for TFAP2A-dependent transcription in regulating amnion fate progression (Sekulovski et al., 2024). However, changes in cellular differentiation states during amnion lineage progression remain largely unexplored.

In this study, we used single cell RNA sequencing (scRNA-seq) analysis to examine the dynamics in gene expression that accompany amnion differentiation of hPSC (herein referred to as the hPSC-amnion) cultured in a soft gel environment, called Gel-3D (Shao et al., 2017a). While no exogenous BMP is added in this Gel-3D amnion model, BMP signaling is activated in the cells by a mechanosensitive cue provided by the soft substrate, thereby initiating human amniogenesis (Shao and Fu, 2022; Shao et al., 2017a; Taniguchi et al., 2019). Interestingly, our data reveal contiguous amniogenic cell states: pluripotency-exiting, early progenitor, late progenitor, specified and maturing, each of which shows transcriptional similarities to distinct cell types in a Carnegie stage 7 human embryo (Tyser et al., 2021). Our data also reveal the presence of two non-amniotic cell types in the Gel-3D system: primordial germ cell-like and advanced mesoderm-like cells. Importantly, we identify a cohort of markers specific to each of these amnion lineage progressing states, and validate selected markers for their expression in the amnion from cynomolgus macaque (*Macaca fascicularis*) embryos. Strikingly, we show that CLDN10, a marker of the progenitor population, exhibits a restricted amniotic expression pattern at the boundary between the amnion and the epiblast of the cynomolgus macaque peri-gastrula. This further supports our findings from a separate study (Sekulovski et al., 2024), which show that the amnion-epiblast boundary is a site of active amniogenesis in the macaque peri-gastrula. Furthermore, loss of CLDN10 results in the formation of primordial germ cell-like cells at the expense of amnion cells. Together, this study provides important single cell-level insight into human amnion fate progression and presents additional evidence for the presence of amnion progenitor cells in primate embryos undergoing gastrulation.

## RESULTS

Previously, we described culture conditions (Gel-3D) in which plated hPSC form polarized cysts initially composed of columnar pluripotent cells, which then spontaneously undergo squamous morphogenesis and begin to express amnion markers (Shao et al., 2017a). Specifically, singly dissociated hPSC are plated densely on a soft gel substrate to form aggregates and are allowed to grow for 24 hours (hrs) in the presence of a ROCK inhibitor (Y-27632). After the first 24 hrs (day 1, d1), the ROCK inhibitor is removed (a trigger to initiate lumenal cyst formation (Hamed et al., 2023; Shao et al., 2017a; Taniguchi et al., 2015)), and a diluted extracellular matrix gel overlay (2% v/v, e.g., Geltrex) is added to provide an additional 3D cue. Immunofluorescent (IF) staining at this time shows cuboidal cells surrounding a central lumen; these cells express NANOG, but not TFAP2A (**Fig. 1A**).

**Figure 1.**
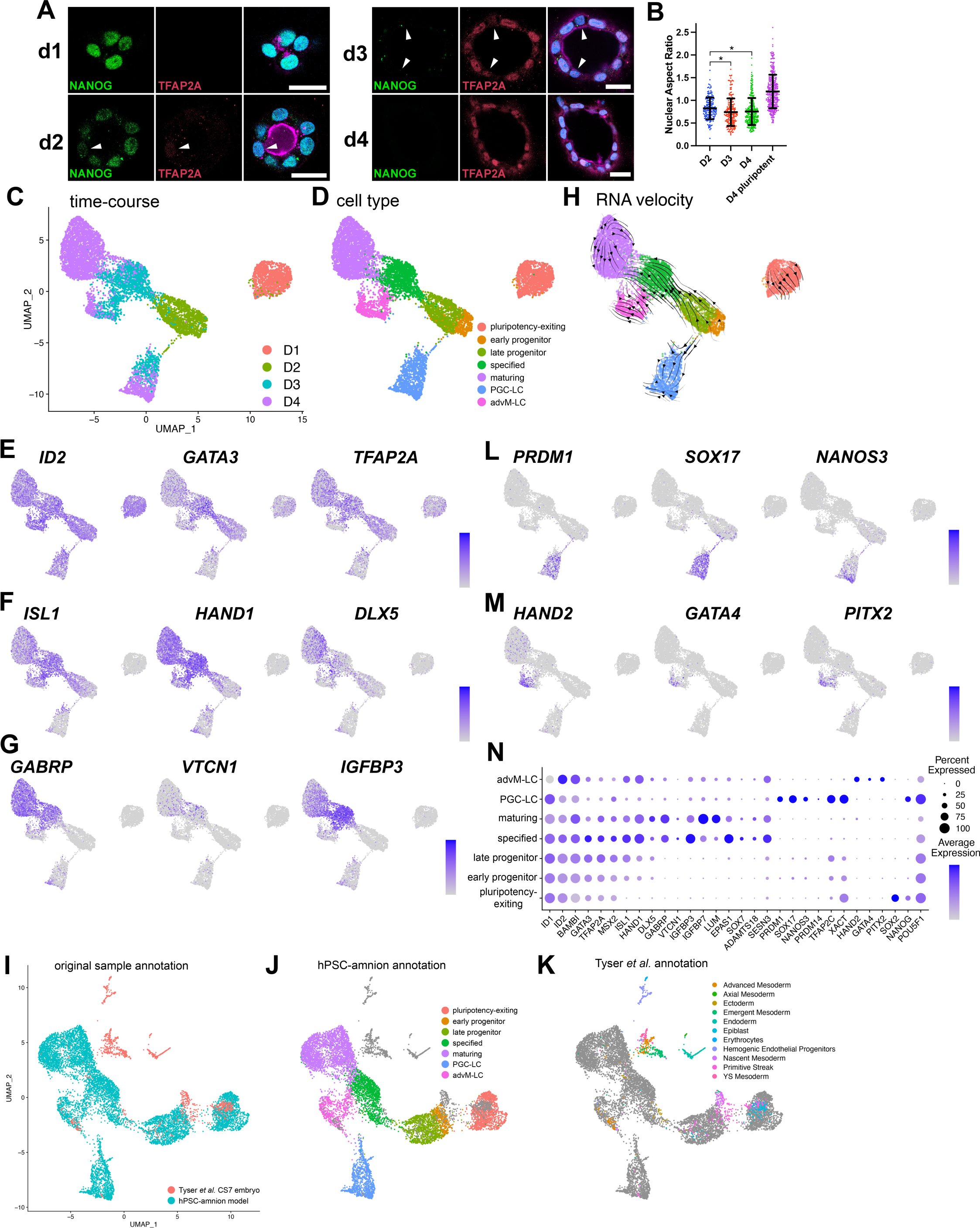
Single-cell transcriptomic signatures of developing Gel-3D human amnion model. (A) Confocal images of d1, d2, d3 and d4 hPSC-amnion cysts stained for TFAP2A, an amnion marker, NANOG, a pluripotency marker, as well as for DNA (blue) and membrane (magenta, using wheat germ agglutinin). White arrowheads indicate cells that display weak TFAP2A expression. Scale bars = 20μm. (B) Quantitation for nuclear aspect ratio, a measure of epithelial cell morphology, of developing hPSC-amnion at indicated timepoints, revealing a gradual squamous morphogenesis (quantitation method in **Fig. S1A**). * indicates p < 0.05. (C,D) A UMAP plot displaying the single cell transcriptomes of 1,359 d1 (salmon), 1,546 d2 (sage), 2,352 d3 (light blue) and 3,508 d4 Gel-3D cells (C) with seven identified cell populations (D, pluripotency-exiting (salmon, 1,363 cells), early progenitor (brown, 479 cells), late progenitor (sage, 1,045 cells), specified (green, 1,366 cells) and maturing (purple, 2,795 cells), PGC-LC (primordial germ cell-like cells, blue, 1,055 cells) and advM-LC (advanced mesoderm-like cells, magenta, 662)). (E-G) Expression of known early (E, *ID2*, *GATA3*, *TFAP2A*), specified (F, *ISL1*, *HAND1*, *DLX5*) and late (G, *GABRP*, *VTCN1*, *IGFBP3*) amnion markers superimposed onto the UMAP plot. (H) RNA velocity analysis of the time-course Gel-3D dataset, showing predicted lineage progression trajectories. (I-K) A UMAP plot showing the integrated single cell transcriptomes of d1-d4 Gel-3D and Tyser *et al*. CS7 embryo datasets, revealing close overlap of Gel-3D amnion progressing cells with Tyser Epiblast, Primitive Streak and Ectoderm/Amnion cells (shown in (I) with original sample, (J) with Gel-3D, and (K) with Tyser *et al*. annotations (individual Tyser annotations plotted in **Fig. S1F**)). (L,M) Expression of primordial germ cell (L, *PRDM1*, *SOX17*, *NANOS3*) and advanced mesoderm (M, *HAND2*, *GATA4*, *PITX2*) markers. (N) Summary of marker expression.

Interestingly, 24 hrs after adding the gel overlay (d2), a subset of cells within the cysts displays reduced expression of NANOG, while TFAP2A expression becomes weakly activated (**Fig. 1A**, arrowhead in d2). By d3, NANOG is no longer detectable. Moreover, while many cells exhibit prominent TFAP2A expression, some cells are only weakly positive for TFAP2A (**Fig. 1A**, arrowheads). This pattern of TFAP2A expression is consistent with the previous observation that amniogenesis initiates focally on the pluripotent cysts and then spreads laterally to form fully squamous cysts (**Fig. 1A**, (Sekulovski et al., 2024; Shao et al., 2017a; Shao et al., 2017b)). These molecular changes are accompanied by a decrease in nuclear aspect ratio (a measure of epithelial cell morphology) over time, revealing more flattened (squamous) shapes by d3 and d4 (**Fig. 1B**, quantitation strategy shown in **Fig. S1A**, and previously established in Shao et al., 2017a; Townshend et al., 2020)). These results provide further molecular and structural evidence that the transition from pluripotent to amnion cell types occurs progressively over the cyst, starting from focal initiation sites.

To explore transcriptional programs of the fate transitioning cells during early stages of human amnion development in this model, we performed time-course scRNA-seq analysis of developing hPSC-amnion cysts, harvested at d1, d2, d3 and d4. hPSC-amnion cysts were dissociated into single cells, followed by single cell isolation and labeling using the 10x Genomics Chromium system, cDNA amplification and sequencing followed by demultiplexing (see Material and Methods for additional details). This scRNA-seq dataset was examined using Seurat (Butler et al., 2018) for data filtering, regression for genes associated with cell cycle progression, normalization, variable gene selection and subsequent unsupervised clustering of cells (see Materials and Methods). The data yield seven distinct populations among 8,765 cells (d1 = 1,359 (salmon), d2 = 1,546 (sage), d3 = 2,352 (light blue), d4 = 3,508 cells (purple), **Fig. 1C**), which, after marker and lineage analyses described below, are labeled pluripotency-exiting (salmon color, 1,363 cells), early progenitor (brown, 479 cells), late progenitor (sage, 1,045 cells), specified (green, 1,366 cells) and maturing (purple, 2,795 cells), PGC-LC (primordial germ cell-like cells, blue, 1,055 cells) and advM-LC (advanced mesoderm-like cells, magenta, 662, **Fig. 1D**). Transcriptomic features of these cells are visualized in Uniform Manifold Approximation and Projection (UMAP) plots (**Fig. 1C,D**).

To broadly characterize the transcriptional state of each population, we examined the expression of known amnion markers (Roost et al., 2015; Shao et al., 2017a; Yang et al., 2021; Zhao et al., 2024; Zheng et al., 2022). Consistent with the amniotic lineage, early amnion markers (*ID2*, *GATA3* and *TFAP2A*, **Fig. 1E**, expression superimposed on the UMAP plot in **Fig. 1C**) are widely expressed starting at d1 in pluripotency-exiting cells that also show broad expression of pluripotency markers (**Fig. S1B**), while intermediate markers that label specified amnion (*ISL1*, *HAND1*, *DLX5*, (Sekulovski et al., 2024; Yang et al., 2021; Zhao et al., 2024)) show abundant expression by d3 (**Fig. 1F**). Late amnion genes such as *GABRP*, *VTCN*1 and *IGFBP3* (expressed in the first-trimester human amnion (Roost et al., 2015) and also in other human amnion models (Sekulovski et al., 2024; Shao et al., 2017a; Yang et al., 2021; Zhao et al., 2024; Zheng et al., 2022)) are primarily enriched in the d3 and d4 populations (**Fig. 1G**). Interestingly, most of the d2 cells show the expression of early, but not specified or pluripotency (SOX2, NANOG), genes, suggesting that the d2 cell population may contain transitioning cell types that give rise to specified amnion (brown and sage cells in **Fig. 1D**). Next, RNA-velocity analysis (Bergen et al., 2020) was applied to examine lineage relationships based on the relative abundance of unspliced and spliced mRNA in each cell. Indeed, the majority of the RNA-vector trajectories are directed from the cells in the early progenitor (brown), late progenitor (sage), to specified (green) states (**Fig. 1H**).

In a recent study, we performed a detailed meta-analysis of the Tyser *et al*. scRNA-seq dataset from a human embryo staged at Carnegie stage 7 (CS7) (Tyser et al., 2021) and showed that the rod-shaped cluster of cells annotated as “Ectoderm” is comprised of lineage progressing, lineage committed, and fully differentiated amnion cell types (Sekulovski et al., 2024). The Tyser *et al*. UMAP plot is reproduced in **Fig. S1C** with the original coordinates and annotations (expression of late markers, *GABRP*, *VTCN*1 and *IGFBP3*, found at the distal tip of the rod-shaped “Ectoderm” population, **Fig. S1C-E**). To examine which cells in the CS7 human embryo share transcriptomic similarities to amnion progressing cells in Gel-3D, the scRNA-seq datasets from the Tyser *et al*. CS7 human embryo and our d1-d4 Gel-3D time-course samples were combined and normalized using an integration feature based on canonical correlation analysis in Seurat (Hao et al., 2021) (**Fig. 1I-K**). Although the combined UMAP plot of this integrated dataset shows some changes in shape from our original UMAP, it is clear that “Epiblast”, “Primitive Streak” and “Ectoderm” populations in the Tyser *et al*. dataset closely map to the amnion progressing populations in our time-course Gel-3D dataset (**Fig. 1I**, see **Fig. S1F** for mapping of each population). Indeed, the amniotic Tyser “Ectoderm” cells primarily overlap with Gel-3D “specified” and “maturing” cell populations, consistent with their amniotic fate; most Tyser “Epiblast” cells are seen in the “pluripotency-exiting” population (**Fig. 1J,K**, **Fig. S1F**). Interestingly, Tyser “Primitive Streak” cells are populated across several Gel-3D amniogenic cell states (**Fig. 1J,K**, **Fig. S1F**), overlapping with the early (brown) and late (sage) progenitor populations, suggesting that some Tyser “Primitive Streak” cells may be actively transitioning, and that our early and late progenitor cells may display transcriptomic characteristics of fate transitioning cells in human peri-gastrula.

Additionally, we identify two non-amniotic cell types that show transcriptional characteristics of PGC- (blue) and advM- (magenta) like cells (**Fig. 1D**). The Tyser-Gel-3D integration analysis shows that the blue cells overlap with the Tyser “Primitive Streak” cells that abundantly express PGC markers (e.g., *PRDM1*, *SOX17*, *NANOS3*, *PRDM14*, *NANOG*, *XACT*, **Fig. 1L**). Several Tyser “Advanced Mesoderm” cells are seen in the magenta cells, and our marker analysis shows that developed mesoderm makers (*HAND2*, *GATA4*, *PITX2*, *ISL1*, *HAND1*) are enriched (**Fig. 1M**, summary of additional markers found in **Fig. 1N**). Strikingly, the RNA velocity trajectories are directed from the interface of the early/late amnion progenitor populations to the PGC-LC (**Fig. 1H**), suggesting their close transcriptomic characteristics. These findings support a growing notion that amnion and PGC progressing cells initially share a common intermediate lineage (Castillo-Venzor et al., 2023; Chen et al., 2019; Xiao et al., 2024; Zheng et al., 2022). The advM-LC population may have formed due to high BMP signaling (Schultheiss et al., 1997; Tsaytler et al., 2023; van Wijk et al., 2007).

To identify genes that are unique to each cluster, unsupervised differential gene enrichment analysis was performed (**Table 1**, adjusted p-value < 0.05). As expected, pluripotency markers are seen in the pluripotency-exiting cluster, and the specified and maturing clusters show enriched expression of known late amnion markers as well as previously uncharacterized genes (*ADAMTS18*, *SESN3* (**Fig. 1N**) – RNA *in situ* hybridization in a cynomolgus macaque (*Macaca fascicularis*) embryo staged between CS12 and CS13, an organogenesis stage, shown in **Fig. S2A-C**).

**Table 1.** Lists of differentially expressed genes in pluripotency-exiting (1506), early progenitor (692), late progenitor (697), specified (944), maturing (868), PGC-LC (1181) and advM-LC (746) clusters.

The most differentially expressed genes in early (brown) and late (sage) progenitor populations are *TBXT* (T-box transcription factor T) and *CLDN10* (a gene encoding Claudin-10, a component of the tight junction), respectively (**Fig. 2A,B**, top three genes plotted in **Fig. 2C**). Recently, we showed that amnion lineage progression in Glass-3D^+BMP^, another model of human amniogenesis, traverses an intermediate transcriptional phase weakly expressing *TBXT* (*TBXT*^low^) before specified markers (e.g., *ISL1*, *HAND1*, *DLX5*) are expressed, revealing lineage progressing characteristics of the cells in the *TBXT*^low^ transcriptional phase (Sekulovski et al., 2024). The same study also showed that cells displaying transcriptomic characteristics consistent with fate progression from *TBXT*^low^ to *ISL1^+^*/*HAND1^+^* specified state are also seen in the Tyser *et al*. CS7 human embryo as well in the Yang *et al*. d14 cynomolgus macaque embryo datasets (Sekulovski et al., 2024). In Gel-3D, while most cells in the early progenitor population express some level of *TBXT*, only a few cells express abundant *TBXT* (**Fig. 2A**). Indeed, our IF time-course analysis shows that cells with weak TBXT expression are seen at d2 in several cells that are not yet expressing ISL1 (**Fig. 2D**, arrowheads); TBXT expression is largely missing in d3 and d4 cysts (**Fig. 2D**), confirming the presence of an early and transient *TBXT*^low^ state in developing Gel-3D hPSC-amnion cysts.

**Figure 2.**
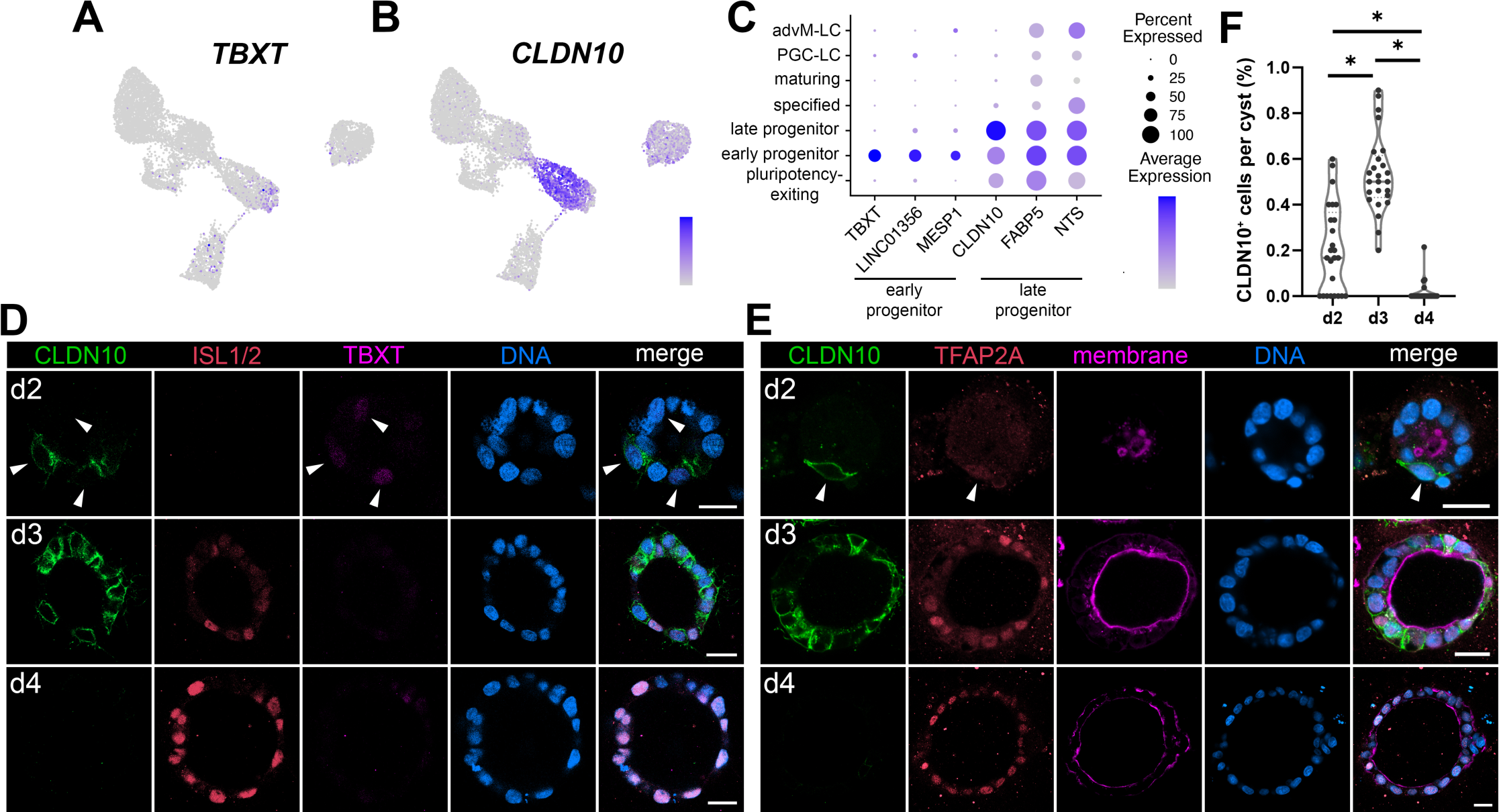
Transient expression of TBXT and CLDN10 marks amnion progenitor populations. (A,B) Expression of *TBXT* (A) and *CLDN10* (B) superimposed onto the time-course Gel-3D UMAP plot. (C) Expression summary of top three most differentially expressed genes in early progenitor and late progenitor populations. (D,E) Confocal micrographs of developing hPSC-amnion harvested at indicated timepoints, stained with indicated markers. (D) Weak TBXT expression is present at d2 (white arrowheads), but is diminished by d3. (E) A white arrowhead indicates a CLDN10/TFAP2A double-positive cell. CLDN10 expression is extinguished by d4. Membrane was stained using wheat germ agglutinin. Scale bars = 20μm (F) Quantitation of CLDN10 expressing cells per cyst in Gel-3D overtime (n = 25 cysts per timepoint).

Prominent *CLDN10* expression is seen throughout the late progenitor population (*CLDN10*^high^), but is rapidly diminished in the specified cells (**Fig. 2B**). Although at lower levels, some *CLDN10* expression is observed in the pluripotency-exiting and early progenitor states (*CLDN10*^low^, **Fig. 2B,C**), consistent with our previous bulk-RNA sequencing analysis showing that low *CLDN10* expression is seen in pluripotent cells (Shao et al., 2017a). Using IF staining in Gel-3D hPSC-amnion model, we validated a dynamic expression pattern of CLDN10 over time (**Fig. 2D,E**). Despite broadly expressed at transcript levels in hPSC-amnion cysts at d2 (**Fig. 2B**), CLDN10 expression is seen in the small fraction of cells that are weakly positive for TBXT and TFAP2A (**Fig. 2D,E**, arrowheads, quantitation in **Fig. 2F**). At d3, CLDN10 is co-expressed with TFAP2A in several, but not all cells (**Fig. 2E**). Further, ISL1^high^ cells do not show clear CLDN10 membrane staining (**Fig. 2D**). In contrast, most cells at d4 retain abundant TFAP2A and ISL1 expression but do not express CLDN10 (**Fig. 2D,E**). Importantly, our analysis for nuclear aspect ratio (NAR) shows that, at d2, CLDN10^+^ cells (mean NAR = 0.54 ± 0.17 STDEV, n = 39) are significantly more squamous compared to CLDN10^-^ cells (NAR = 0.97 ± 0.31 STDEV, n = 87, p<0.001). While previous studies implicated *CLDN10* as an amnion marker (Zheng et al., 2019; Zheng et al., 2022), a detailed analysis was not performed. Thus, these data present evidence suggesting that CLDN10^high^ marks a later transient progenitor state that follows an earlier *TBXT*^low^/*CLDN10*^low^ progenitor state, but CLDN10 expression is extinguished in more differentiated cells. Single cell transcriptomic characteristics consistent with this lineage progression from the *TBXT*^low^/*CLDN10*^low^ state to the *CLDN10*^high^ state is also seen in the Tyser *et al*. CS7 human embryo as well in the Yang *et al*. d14 cynomolgus macaque embryo datasets (**Fig. S3A-C**).

Interestingly, in PASE (post-implantation amniotic sac embryoids), an *in vitro* model of human amniotic sac formation (Shao et al., 2017b), CLDN10 is localized specifically to cells at the boundary between squamous amnion and columnar pluripotent cells (**Fig. 3A**, arrowheads). To test whether CLDN10^+^ progenitor-like boundary cells are present in the developing primate amniotic sac *in vivo*, we analyzed CLDN10 expression in a cynomolgus macaque embryo displaying the amniotic sac (**Fig. 3B**, staged between CS6 and CS7). Strikingly, CLDN10 expression is exclusively seen at the boundary separating amnion and epiblast tissues. These results are consistent with our recent findings indicating that this boundary is likely a site of active amniogenesis (i.e., epiblast cells at the boundary actively undergo amnion specification) in the primate peri-gastrula (Sekulovski et al., 2024). Importantly, a cynomolgus macaque embryo from a later stage (**Fig. 3C,D**, CS10) also shows CLDN10 positive cells at the amnion-embryo boundary. In the CS10 embryo, while a collection of cells expressing weak CLDN10 is seen posteriorly (**Fig. 3C,D** – inset (iii)), CLDN10 expression is highly prominent anteriorly in the cells at the boundary of the amnion and the developing mediolateral placode (**Fig. 3C,D** – inset (i), additional images in **Fig. S3D,E**). Together, these results establish the presence of previously unrecognized CLDN10^+^ amnion-embryo boundary population that primarily gives rise to amnion during gastrulation as well as during early organogenesis in the cynomolgus macaque.

**Figure 3.**
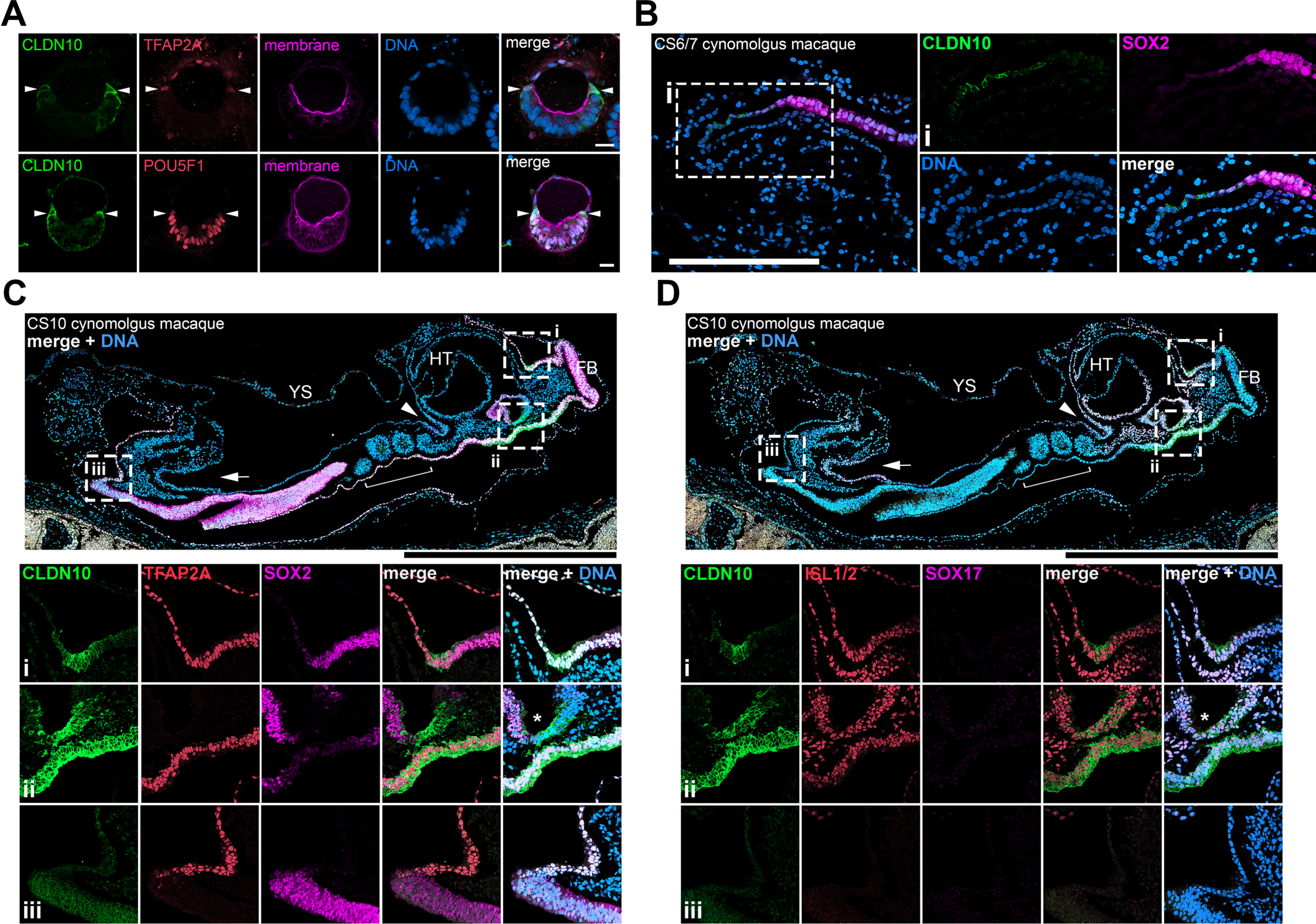
CLDN10 is expressed in amnion progenitor cells at the amnion-epiblast boundary of the PASE model as well as cynomolgus macaque peri-gastrula. (A) Optical sections of PASE stained with indicated markers. White arrowheads indicate CLDN10 staining at the amnion-epiblast boundary. Weak CLDN10 signal in the pluripotent territory is likely due to weak CLDN10 expression in the pluripotent cells; brighter signal in the pluripotent territory is likely caused by the boundary cell extensions. (B) Confocal images of a cynomolgus macaque embryo staged at CS6/7 stained using indicated markers. CLDN10 is expressed at the transitioning boundary cells between the amnion and the epiblast. Insets indicate magnified regions. (C,D) Confocal images of cynomolgus macaque embryos staged at CS10 stained using indicated markers. Insets indicate magnified regions. In the CS10 embryo (early organogenesis stage), while some posterior boundary CLDN10 staining is seen (seen in (iii)), the anterior CLDN10 staining is highly prominent at the amnion-anterior surface ectoderm border (in (i)). CLDN10 is also enriched in the dorsal foregut labeled by ISL1/2 and SOX2 but not SOX17 (indicated by * in (ii)). **Fig. S3F** provides a bird’s-eye view of the CS10 embryo. HT, heat tube; FB, forebrain; YS, yolk sac. White arrowheads indicate the foregut pocket, while white arrows indicate the hindgut pocket. Scale bars = 20μm (A), 200μm (B), 500μm (C,D).

To investigate the role of CLDN10, H9 hESC lacking CLDN10 were cultured in the Gel-3D condition (**Fig. 4A-D**, see **Fig. S4** for generation of *CLDN10*-KO lines). Morphologically, while *CLDN10*-KO cyst formation is largely similar to controls by d3, by d4, squamous morphogenesis as well as overall cyst formation are disrupted. Interestingly, our IF analysis shows that, in the KO background, SOX2 expression gradually reduces overtime similar to controls, but cells expressing ISL1 are reduced by d4 (**Fig. 4A,B**). Given the presence of PGC-LC in Gel-3D (**Fig. 1D,L**), we next examined the expression of PGC markers. Strikingly, while very few cells are NANOG^+^/SOX17^+^ PGC-LC in controls at d4, NANOG/SOX17 double positive cells are seen prematurely at d2 as well as at d3 (**Fig. 4C,D**). These results suggest that CLDN10 functions to promote amnion fate progression, and prevent emergence of the PGC-like lineage.

**Figure 4.**
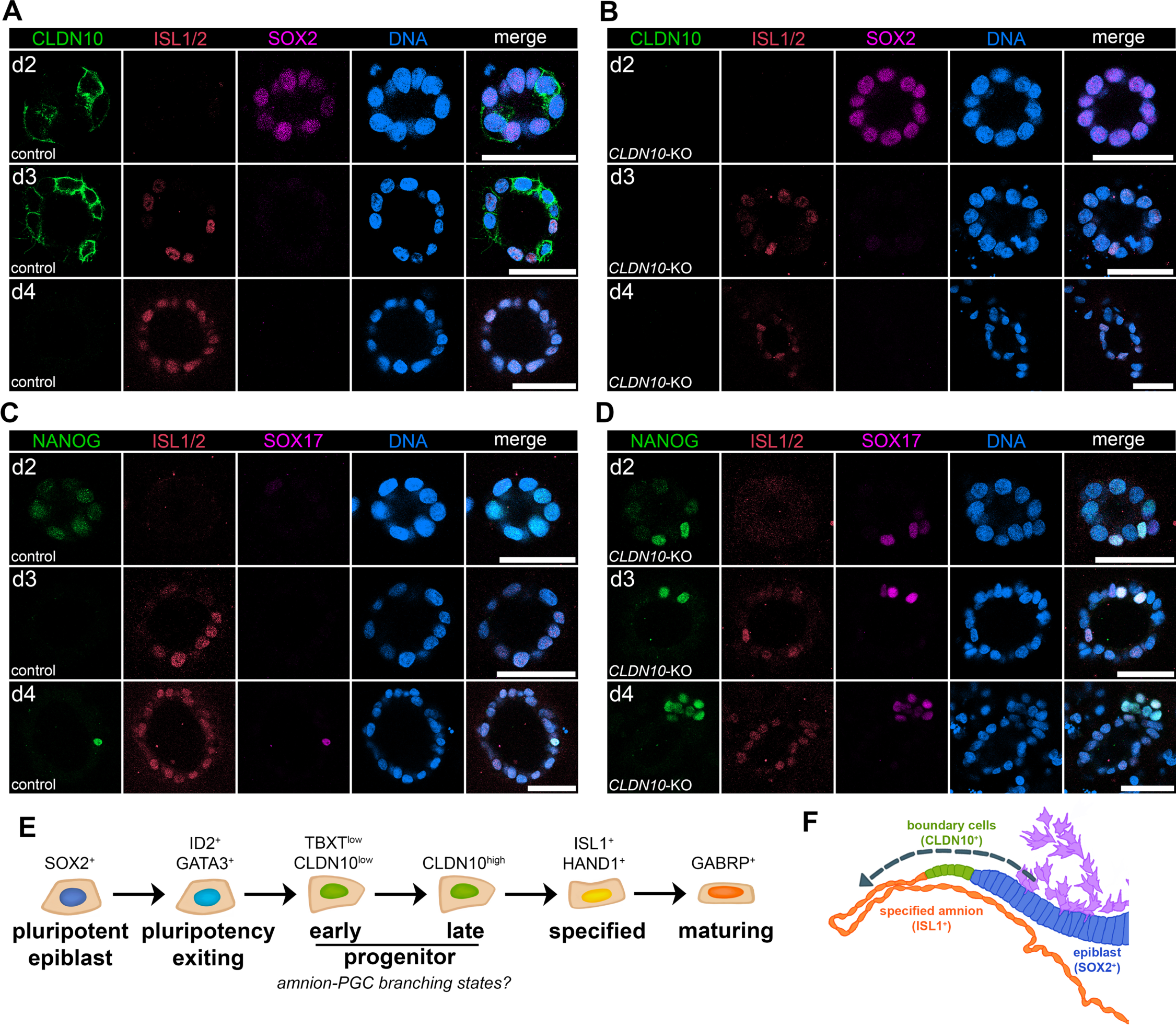
CLDN10 is critical for maintaining amnion fate progression in the progenitor population. (A-D) Confocal images of control (A,C) and *CLDN10*-KO (B,D) hPSC grown in Gel-3D, harvested at indicated timepoints and stained with indicated markers. CLDN10 expression is not seen in the KO (B). In the absence of *CLDN10*, PGC-LC formation is seen early in d2 cysts, and cyst organization is disrupted by d4. Scale bars = 50μm. (E) Flow chart outlining five continuous amniogenic lineage progressing states with representative markers. (F) A schematic representation of primate peri-gastrula near the primitive streak (generated based on images in Fig. 3B, purple cells indicate primitive streak-derived disseminating cells). Transcriptional signatures consistent with amniotic lineage progression (indicated by dotted arrow) are observed at the amnion-epiblast boundary: progressing from SOX2^+^ epiblast cells (blue), CLDN10^+^ transitioning boundary cells (green), and, then, to ISL1^+^ lineage specified amnion (orange).

## DISCUSSION

Together, this study presents a temporally resolved single cell transcriptomic resource for future investigations of the amniogenic transcriptional cascade in developing primate embryos. In summary, we have profiled the transcriptomic signatures of lineage progressing cells in an *in vitro* model of hPSC-amnion development, and identified 1) five contiguous amnion progressing states (outlined in **Fig. 4E**) including two previously unrecognized amnion progenitor populations, 2) an amnion progenitor population labeled by CLDN10 at the boundary between the amnion and the epiblast of the primate amniotic sac (model schematic in **Fig. 4F**), and 3) the role of CLDN10 in maintaining amnion fate progression.

Our time-course scRNA-seq analysis showed that PGC-LC are present in our Gel-3D model, and that some of the amnion progenitor population may contribute to PGC-LC, supporting a growing notion that amnion and PGC progressing cells initially share a common lineage (Castillo-Venzor et al., 2023; Chen et al., 2019; Xiao et al., 2024; Zheng et al., 2022), and providing detailed transcriptomic insights into this lineage diversification step that occurs in the Gel-3D system (**Fig. 4E**). Moreover, the loss of CLDN10, a marker of the late amnion progenitor and a tight junction component, leads to increased PGC-LC formation, while amnion formation is reduced. Interestingly, a recent study by Vasic *et al*. showed that reduced level of TJP1 (also known as ZO-1) in unconfined hPSC monolayer colonies treated with BMP4 (a system that undergoes gastrulation-like patterning (Joy et al., 2021) similar to the micropatterned gastruloid model, (Heemskerk et al., 2017; Warmflash et al., 2014) leads to increased emergence of a germ cell lineage (Vasic et al., 2023). Therefore, our results present additional evidence for the role of tight junction formation in suppressing PGC lineages, and demonstrate a key role of CLDN10 in maintaining amnion fate progression in progenitor cells. Given that Claudin proteins control tight junction properties and epithelial characteristics (Angelow et al., 2008), CLDN10 might also be critical for organizing a boundary structure containing shape/fate-transitioning cells flanked by squamous amnion or columnar/pseudostratified pluripotent cells on each side.

Previous studies established that BMP is a major trigger of amniogenesis. What other potential mechanisms could help to initially induce and maintain the half amnion-half epiblast structure? Trophectoderm may play some roles in triggering amnion specification given that these cells directly overlay the nascent amnion during implantation. Indeed, molecular characterization of early cynomolgus macaque embryos by Sasaki *et al*. (Sasaki et al., 2016) have suggested that the trophectoderm may provide some cue (e.g., secreted ligands) to the underlying epiblast cells. Also, a recent study by Pedroza *et al*. showed that co-culturing hPSC on a monolayer of human trophoblast stem cells helps initiate amniogenesis (Pedroza et al., 2023).

Furthermore, visceral endoderm might play an important role in defining the zone in which amniogenesis can occur, helping to shape the amnion-epiblast boundary territory. Studies have established that visceral endoderm plays a major role in embryo patterning by expressing secreted BMP antagonists (Chambers et al., 2009; Perea-Gomez et al., 2002; Shawlot et al., 1999), even at very early stages of implantation (Bergmann et al., 2022; Sasaki et al., 2016). Therefore, it is possible that, during implantation, epiblast cells that are in close proximity to visceral endoderm contribute to embryo proper because the presence of secreted BMP antagonists prevents them from undergoing amniogenesis. However, more distal epiblast cells at the uterine-proximal pole of the epiblast cyst, which are farther away from the BMP antagonist source, can respond to BMP signaling and form amnion. The amnion-epiblast boundary territory might be the most distal region at which amniogenesis can take place, and the balance between the rate of epiblast proliferation and the rate of amniogenesis at the boundary likely contributes to maintaining the ratio between amnion and epiblast cells in implanting embryos, as well as in peri-gastrula.

Recently, single cell transcriptomics analysis has emerged as a valuable tool to ground findings from *in vitro* models with natural as well as cultured primate embryo systems (e.g., (Yang et al., 2021; Zhao et al., 2024; Zheng et al., 2022)). To aid in additional investigations, several culture systems have been developed to generate amnion, and single cell transcriptomic analyses have been performed in several of the systems (e.g., (Chen et al., 2019; Minn et al., 2020; Overeem et al., 2023; Rostovskaya et al., 2022)). Future transcriptomic comparisons of the amnion in each of these models will enable us to gain additional insights into amniogenic mechanisms.

## MATERIALS AND METHODS

### hESC lines

Human embryonic stem cell line H9 was used in this study (WA09, P30, P48, WiCell; NIH registration number: 0062). All protocols for the use of hPSC lines were approved by the Human Stem Cell Research Oversight Committee at the Medical College of Wisconsin and the Human Pluripotent Stem Cell Research Oversight Committee at the University of Michigan. All hPSC lines were maintained in a feeder-free system for at least 20 passages and authenticated as karyotypically normal at the indicated passage number. Karyotype analysis was performed at Cell Line Genetics. All hPSC lines tested negative for mycoplasma contamination (LookOut Mycoplasma PCR Detection Kit, Sigma-Aldrich). In summary, hESC were maintained in a feeder-free culture system with mTeSR1 medium, or with 50%/50% mix of mTeSR1 and mTeSR plus (STEMCELL Technologies). hESC were cultured on 1% (v/v) Geltrex (Thermo Fisher Scientific), or with Cultrex SCQ (Bio-techne) coated 6 well plates (Nunc). Cells were passaged as small clumps every 4 to 5 days with Dispase (Gibco). All cells were cultured at 37°C with 5% CO_2_. Media was changed every day. hESC were visually checked every day to ensure the absence of spontaneously differentiated, mesenchyme-like cells in the culture. Minor differentiated cells were scratched off the plate under a dissecting scope once identified. The quality of all hESC lines was periodically examined by immunostaining for pluripotency markers and successful differentiation to three germ layer cells. Similar methods were previously used in (Sekulovski et al., 2024; Wang et al., 2021).

### Gel-3D hPSC-amnion formation assays

Methods for these assays have been previously described (Shao et al., 2017a; Shao et al., 2017b) with exceptions that 60-70uL of undiluted ECM gel solution is hand-streaked on ice cold 22mmx22mm square coverslips, and that, in addition to Geltrex (Life Technologies), Cultrex Ultimatrix (Bio-Techne) was also used. Singly dissociated cells were prepared using Accutase (Sigma-Aldrich) and were plated on ECM gel-coated coverslips at 25,000 cells/cm^2^ in mTeSR1 or in 50%/50% mTeSR1/mTeSR plus mix medium in the presence of 10µM Y-27632 (STEMCELL Technologies). After 24 hr, cells were then incubated in media containing 2% Geltrex/Ultimatrix overlay without Y-27632 with daily media changes (note that 1% gel overlay was used between d2 to d4).

### DNA constructs

#### piggyBac-CRISPR/Cas9 (pBACON) constructs

A piggyBac-CRISPR/Cas9 (pBACON) vector that contains SpCas9-T2A-puro and hU6-gRNA expression cassettes flanked by piggyBac transposon terminal repeat elements (pBACON-puro), which allows subcloning of annealed oligos containing gRNA sequence at *BbsI* site, has been previously described (Shao et al., 2017b; Townshend et al., 2020; Wang et al., 2021). gRNA targeting genomic sites and oligo sequences to generate pBACON-puro-h*CLDN10*-ICL1 (primers: CRISPR_hCLDN10ICL#1_s and CRISPR_hCLDN10ICL#1_as) are found in **Fig. S4** and **Table 2**; these were designed using a publicly available tool (https://www.idtdna.com/site/order/designtool/index/CRISPR_CUSTOM). Similar methods were previously used in several publications (e.g., (Sekulovski et al., 2024; Wang et al., 2021)).

**Table 2.** List of antibodies, RNAscope probes, primers and plasmids used in this study.

### piggyBac-based transgenic and genome edited hESC lines

PB constructs (3μg) and pCAG-ePBase (Lacoste et al., 2009) (1μg) were co-transfected into H9 hESC (70,000 cells cm^-2^) using GeneJammer transfection reagent (Agilent Technologies). To enrich for successfully transfected cells, drug selection (puromycin, 2μg/mL for 4 days) was performed 48- to 72-hrs after transfection. hESC stably expressing each construct maintained the expression of pluripotency markers.

During pBACON-based genome editing, puro-selected cells were cultured at low density (300 cells cm^-2^) for clonal selection. Established colonies were manually picked and expanded for screening indel mutations using PCR amplification of a region spanning the targeted gRNA region (genomic DNA isolated using DirectPCR Lysis Reagent (Tail) (VIAGEN), primers: hCLDN10, Geno_hCLDN10CDS2_RI_fw and Geno_hCLDN10CDS2_NI_rv), which were subcloned into pPBCAG-GFP (Chen and LoTurco, 2012) at EcoRI and NotI sites, and sequenced (Seq-3’TR-pPB-Fw). At least 12 to 15 bacterial colonies were sequenced (Sanger sequencing) to confirm genotypic clonality. Control cells are wild-type, unedited H9 hESC in all loss-of-function experiments.

### Cynomolgus macaque

#### Animals

The female and male cynomolgus macaques were housed and cared for at the Wisconsin National Primate Research Center (WNPRC). All procedures were performed in accordance with the NIH Guide for the Care and Use of Laboratory Animals and under approval of the University of Wisconsin College of Letters and Sciences and Vice Chancellor Office for Research and Graduate Education Institutional Animal Care and Use Committee (protocol g005061).

#### Animal breeding and pregnancy detection

Beginning on day 8 post-onset of menses, the female was housed with a compatible male and monitored for breeding. Animals were pair-housed until day 16-20 post-onset of menses. A 2-3 mL blood draw was performed daily from day 8 post-onset of menses until day 16 to assess the timing of ovulation based on the estradiol peak and rise in progesterone in serum. Serum samples were analyzed for estradiol (25 μL) and progesterone (20 µL) using a cobas e411 analyzer equipped with ElectroChemiLuminescence technology (Roche, Basal, Switzerland) according to manufacturer instructions. Results were determined via a calibration curve which was instrument-generated by 2-point calibration using traceable standards and a master curve provided via the reagent barcode. Inter-assay coefficient of variation (CV) was determined by assaying aliquots of a pool of rhesus plasma. For estradiol, the limit of quantitation (LOQ) was 25 pg/mL, the intra-assay CV was 2.02%, and the inter-assay CV was 5.05%. For progesterone, the LOQ was 0.2 ng/mL, the intra-assay CV was 1.37%, and the inter-assay CV was 4.63%. A transabdominal ultrasound was performed to detect pregnancy as early as 14 days post-ovulation. The ultrasound measurements in combination with the timing of ovulation were used to estimate the day of conception and gestational age of the pregnancy.

#### Terminal perfusion uterine collection, paraffin embedding, sectioning and staining

The pregnant females were sedated with intramuscular ketamine (>15 mg/kg) followed by IV sodium pentobarbital (>35 mg/kg) and then perfused with 4% paraformaldehyde (PFA) via the left ventricle. The entire uterus and cervix were removed. The serosa and superficial myometrium were scored for better fixative penetration and to denote dorsal and ventral orientation. Tissues were fixed in 4% PFA with constant stirring and solution changes were performed every 24 hrs for a total of 72 hrs. The uterus was serially sectioned from the fundus to the internal cervical os into 4 mm slices using a dissection box. Cassettes with tissue sections were transferred into 70% ethanol, routinely processed in an automated tissue processor and embedded in paraffin for histological analyses (5 µm sections). Fixed uterine tissue were cut in 5 μm thick cross-sections, mounted on slides, deparaffinized in xylene and rehydrated in an ethanol series. Antigen retrieval was performed by boiling in citrate buffer. Sections were blocked 4% goat serum in PBS at RT for at least 3-hr. Subsequent immunolocalization was performed using commercially available primary antibodies, incubated overnight at 4 °C in 4% serum. Immunofluorescent detection was performed using secondary antibodies tagged with a fluorescent dye (fluorophores excitation = 488, 555, and 647nm), and counterstained with DAPI. Negative controls were performed in which the primary antibody was substituted with the same concentration of normal IgG of the appropriate isotype. Images were obtained with a Zeiss LSM980 microscope.

Note that similar methods were used in (Sekulovski et al., 2024).

### Confocal microscopy of fixed samples

Confocal Images of fixed samples were acquired using a Nikon-A1 (Nikon) and a Zeiss LS980 laser scanning confocal microscopes. Non-3D images were generated using Zen, FIJI (NIH) and Photoshop (Adobe).

### Immunostaining

hPSC-amnion cysts grown on the coverslip were rinsed with PBS (Gibco) twice, fixed with 4% paraformaldehyde (Sigma) for 60 min, then rinsed with PBS three times, and permeabilized with 0.1% SDS (Sigma, in 1x PBS) solution for 60 min. The samples were blocked in 4% heat-inactivated goat serum (Gibco) or 4% normal donkey serum (Gibco) in PBS overnight at 4°C. The samples were incubated with primary antibody solution prepared in blocking solution at 4°C for 48 hr, washed three times with PBS (30 min each), and incubated in blocking solution with goat or donkey raised Alexa Fluor-conjugated secondary antibodies (Thermo Fisher), at room temperature for 24 hours. Counter staining was performed using Hoechst 33342 (nucleus, Thermo Fisher Scientific), Alexa-Fluor-conjugated wheat germ agglutinin (membrane, Thermo Fisher Scientific) and Phalloidin (F-ACTIN, Thermo Fisher). All samples were mounted on slides using 90% glycerol (in 1x PBS). When mounting hPSC-cysts samples, modeling clay was used as a spacer between coverslip and slide to preserve lumenal cyst morphology. Antibodies for IF staining are found in **Table 2**. Similar staining method was previously used in (Shao et al., 2017a; Shao et al., 2017b).

### RNA isolation and Single cell RNA sequencing

Gel-3D hPSC-amnion samples were rinsed once with cold 1x PBS prior to treating with Cultrex Organoid Harvesting solution (R&D Systems) to non-enzymatically depolymerize Ultimatrix (gel bed, R&D systems), which were then dissociated into single cell suspensions using the Neural Tissue Dissociation Kit (P) (Miltenyi Biotech, 130-092-628) following the manufacturer’s protocol with the exception that gentle agitations were performed manually with a P1000 pipette and all incubations were performed at 10°C. Cells were filtered through 70μm cell strainers (SP Bel-Art, 136800070), rinsed with ice cold 1x PBS and counted before proceeding with the 10x Chromium Next Single Cell 3’ v3.1 platform (single cell preparation, single cell isolation and barcoding performed according to the manufacturer’s instructions). Libraries were sequenced (Illumina NovaSeq 6000 S4 flowcell, the DNA services laboratory of the Roy J. Carver Biotechnology Center at the University of Illinois at Urbana-Champaign) followed by quality control (adaptor trimming, de-duplication), alignment and demultiplexing based on manufactured indices using Partek Flow software (Partek Inc. St. Louis, MO, USA, CellRanger).

### Bioinformatics – single cell RNA sequencing dataset analysis

Analysis of the d1-4 time-course amnion scRNA-seq dataset was performed using the Seurat R package (v.4.2.1, (Hao et al., 2021)). For quality control, cells were filtered out if the total number of detected genes was less than 1,500, if the expression percentage from mitochondrial genes was less than 3% or greater than 15%, or if the total number of transcripts was greater than 30,000. Expression values were then log-normalized with scaling factor 10,000 for cell-level normalization, and further centered and scaled across genes. Principal component analysis (PCA) was performed prior to embedding 8,765 cells into two-dimensional space with UMAP using top 15 PCs. Cell clusters were identified using FindNeighbors and FindClusters functions based on a shared nearest neighbor modularity optimization clustering algorithm (resolution set as 0.2). In addition, the “early progenitor” cluster was identified within the original “progenitor” cluster (containing both early and later progenitors) by a gating strategy that sets a cut-off based on the normalized expression levels of TBXT (more than 0.4) and CLDN10 (less than 3.8). Nearby TBXT^-^ cells within the TBXT^+^ domain were also included in the “early progenitor” cluster using CellSelector function because those cells are likely due to dropouts. The cluster marker genes were identified using FindAllMarkers function (Wilcoxon Rank Sum test) with adjusted p values less than 0.05.

#### Mapping and gene expression analyses of Tyser et al. Carnegie Stage 7 human embryo single cell RNA sequencing dataset

The original UMAP coordinates and annotations for all 1,195 cells were used in **Fig. S1C** as well as in the accompanying expression plots. The processed data were downloaded from http://www.human-gastrula.net, which was used to generate a UMAP plot (using DimPlot function in R package Seurat), as well as to perform expression analyses (FeaturePlot function in Seurat).

#### Integration and gene expression analyses of the d1-4 amnion and Tyser et al. Carnegie Stage 7 human embryo single cell RNA sequencing datasets

The two single cell datasets were integrated using a canonical correlation analysis based approach implemented in the IntegrateData function with 4,000 anchor features and 50 dimensions for anchor weighting in Seurat R package (v4.2.1).

#### Integration and gene expression analyses of Yang et al. GD14 cynomolgus macaque embryo single cell RNA sequencing dataset

The processed data were downloaded from GSE148683, and the ensemble genome build Macaca_fascicularis_5.0 release 96 was used to identify human orthogonal gene symbols. Datasets from two distinct GD14 embryos (Yang et al., 2021) were integrated using 5,000 anchor features and 10 dimensions in the IntegrateData function in Seurat R package (v.4.2.1) package: trophoblast cells were removed from the dataset prior to integration. Six general cell populations (epiblast, transition, mesoderm, amnion, endoderm and extraembryonic mesenchyme) were identified using FindClusters function (resolution as 0.4).

#### RNA velocity (scvelo) analysis

From the aligned BAM files, a loom file was generated for each d1-d4 dataset respectively using the function run10x mode in software velocyto (v0.17) with default parameters to create count matrices made of spliced and unspliced read counts (with Human genome annotation hg19). scVelo (Python package v.0.2.5 (Bergen et al., 2020) was then used to analyze the loom files and examine RNA velocity. Inferred RNA velocity was overlaid onto the UMAP embedding (created by Seurat pipeline as described above) and visualized by stream plot (with default parameters except smoothness was set to 1).

---

**Figure S1. Related to Figure 1.**
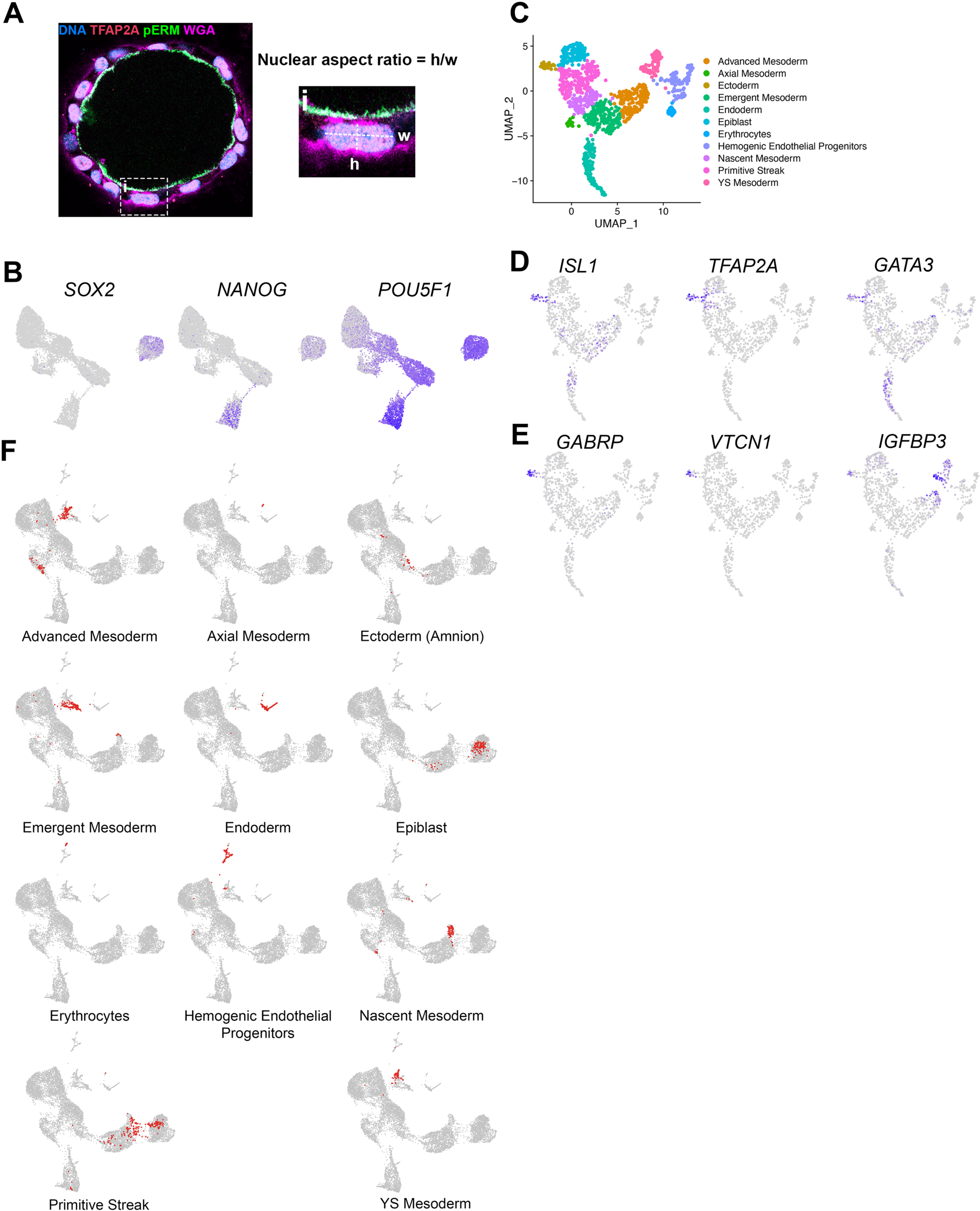
(A) Confocal images of hPSC-amnion stained with indicated markers, outlining nuclear aspect ratio quantitation. (B) Expression of pluripotency markers, SOX2, NANOG and POU5F1, superimposed onto the Gel-3D UMAP plot. (C) A UMAP plot displaying the original single cell transcriptome coordinates of the CS7 human embryo described in Tyser *et al*., shown with the original annotations. (D,E) Expression of indicated broad (D) and late (E) amnion markers superimposed onto the Tyser UMAP plot. (F) UMAP plots showing the integrated single cell transcriptomes of d1-d4 Gel-3D and Tyser *et al*. CS7 embryo datasets with individual Tyser annotations.

**Figure S2.**
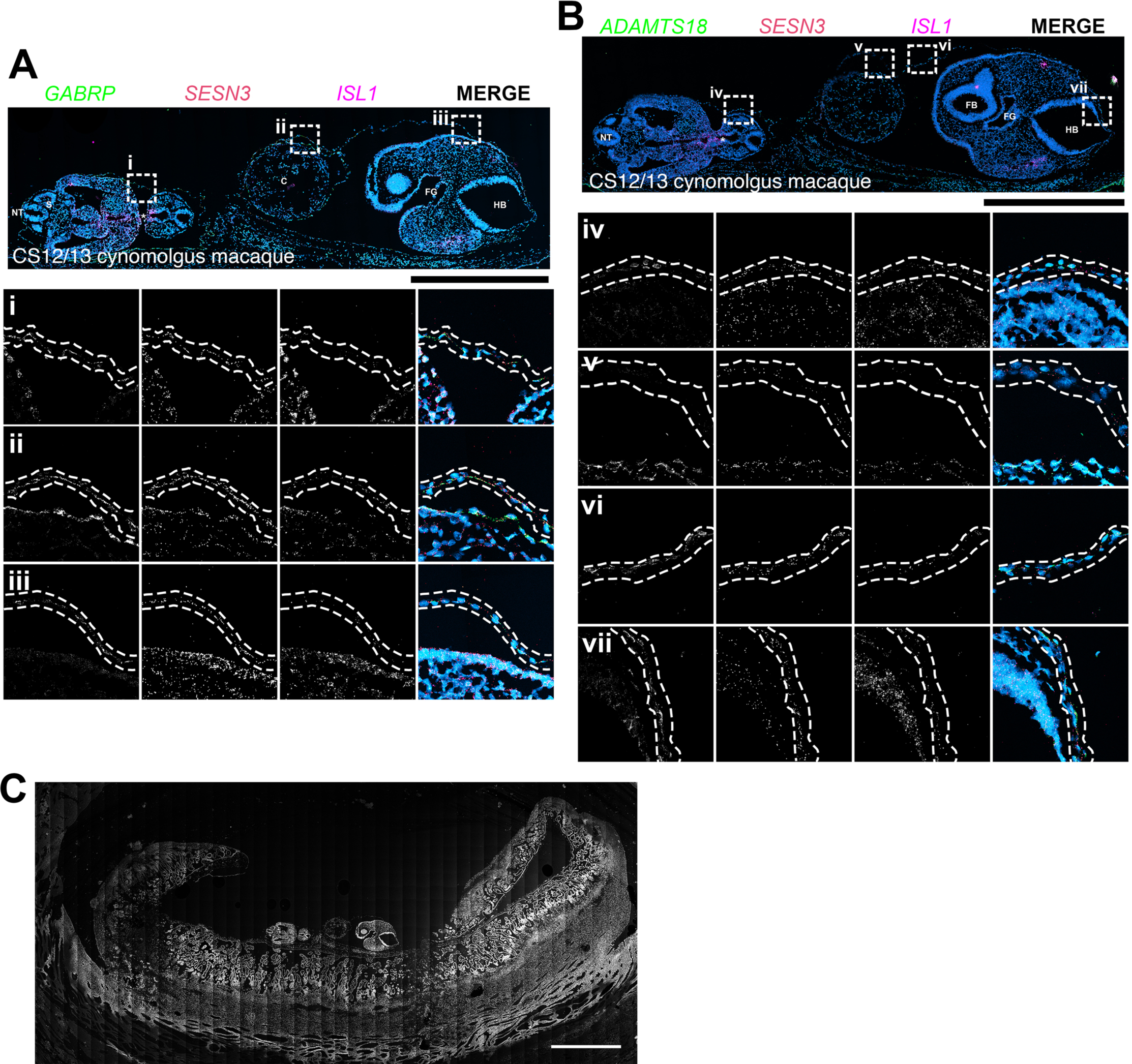
Validation of selected specified and maturing markers in a cynomolgus macaque embryo staged between CS12 and CS13. (A,B) CS12/13 cynomolgus macaque embryo sections were stained for indicated markers (A: *GABRP* – green, *SESN3* – red, *ISL1* – magenta; B: *ADAMTS18* – green, *SESN3* – red, *ISL1* – magenta) using RNA *in situ* hybridization. (i) – (vii) indicate insets: individual channels are shown in gray scale to aid visualization (from left to right: *GABRP*, *SESN3*, *ISL1* and merge in (A), *ADAMTS18*, *SESN3*, *ISL1* in (B)), and dotted lines indicate a layer of amniotic epithelium and mesenchyme. Most amnion cells are positive for *GABRP*, *SESN3* and *ISL1*. *ADAMTS18* expression is restricted to a fraction of cells. In (iv), there are four *ADAMTS18^high^* cells, while *ADAMTS18* is largely undetected in the amnion in (v). In (vi) and (vii), most cells are *ADAMTS18^high^*. FB; forebrain, HB, hindbrain; FG, foregut; C; cardiac tissue; NT, neural tube; S, somite. * indicates an *ISL1^+^* tissue that is likely of genital tubercle lineage. Scale bars = 500μm. (C) An overview image of the CS12/13 embryo implantation site shown in (A). Only nuclear staining is shown in grayscale. Scale bar = 2mm.

**Figure S3.**
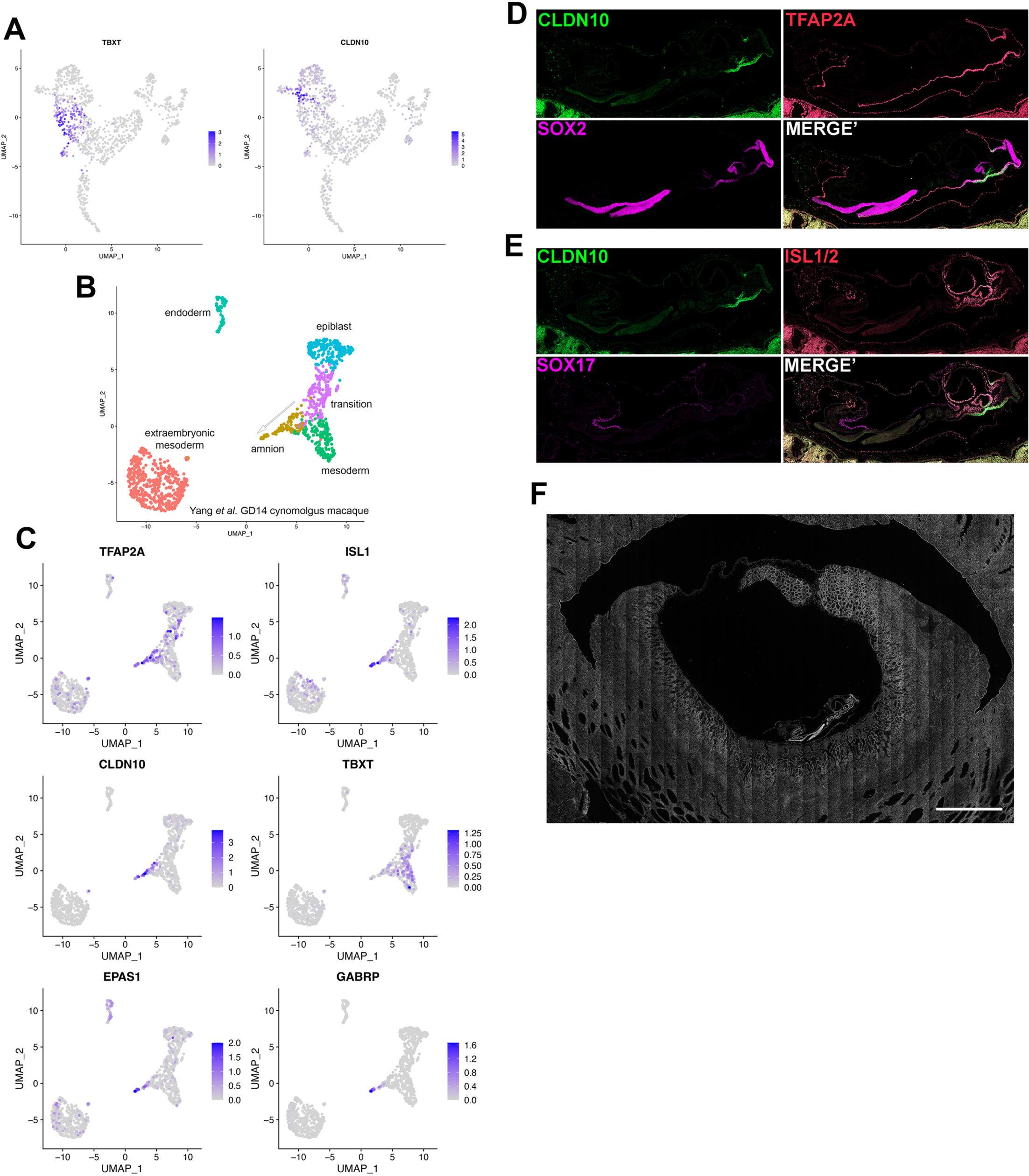
Additional expression analyses. (A) Expression of *TBXT* and *CLDN10* superimposed onto the uncropped Tyser *et al*. CS7 human embryo UMAP plot. (B) A UMAP plot displaying the GD14 cynomolgus macaque single cell transcriptome with six identified general clusters (epiblast – light blue; transition – purple; amnion – gold; mesoderm – green; endoderm – teal; extraembryonic mesoderm – salmon). Arrow indicates the likely trajectory of amnion differentiation based on the data in Yang *et al*. as well as the expression analysis in (C). Note that, although not identical to the published UMAP plot in Yang *et al*., general characteristics are well recapitulated. Trophectoderm cells have been omitted as performed in Yang *et al*.. (C) Expression of indicated markers superimposed onto the Yang *et al*. UMAP plot. Late amnion markers (EPAS1 and GABRP) are expressed at the tip of the TFAP2A and ISL1 double-positive amnion cluster. CLDN10 positive cells show a weak TBXT expression level. (D,E) Individual channels of the whole embryo images in Fig. 3C and 3D, respectively. (F) An overview image of the CS10 (Fig. 3C) embryo implantation sites (DNA signal shown in gray scale, Scale bar = 2mm).

**Figure S4.**
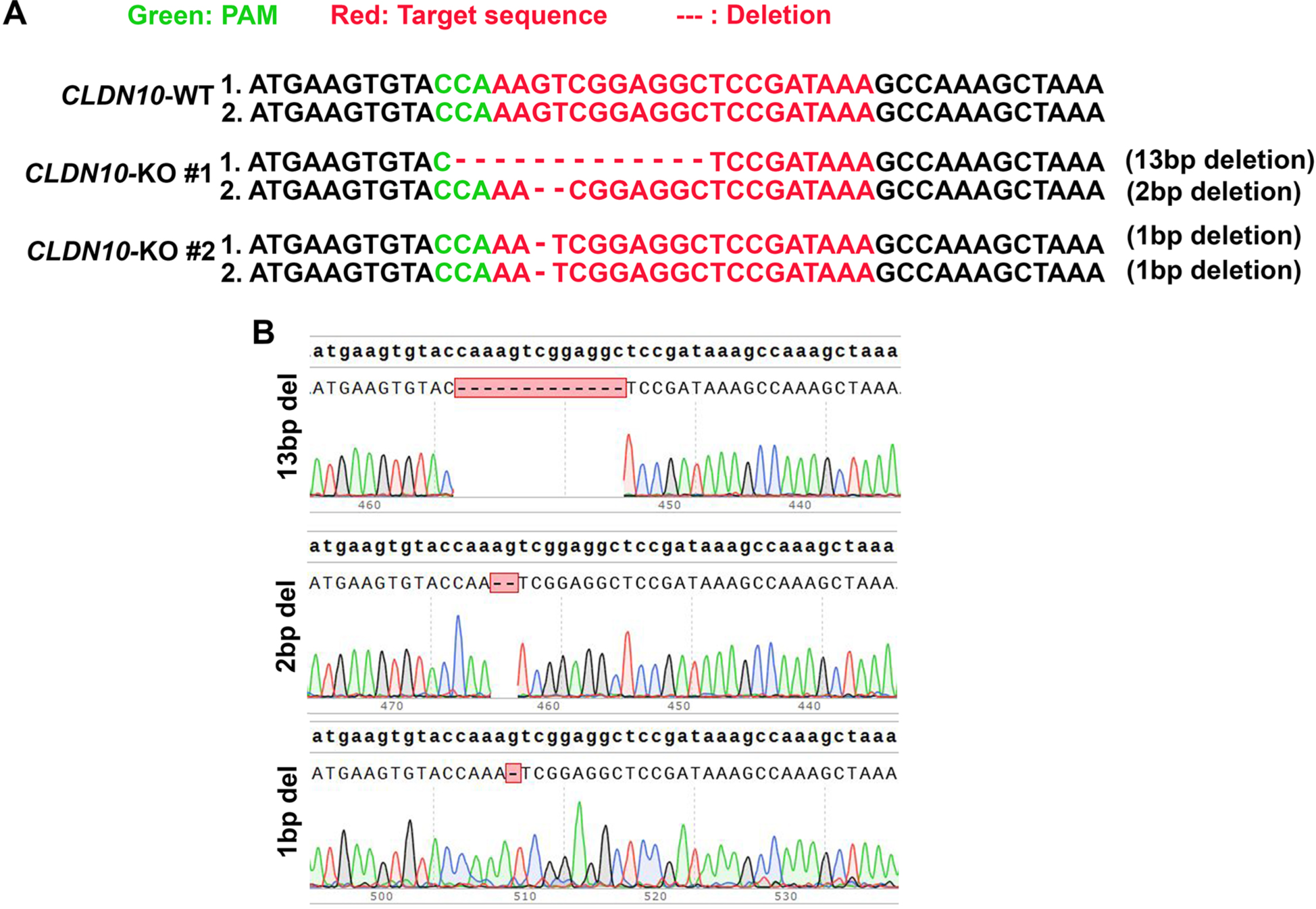
Validation of two distinct *CLDN10*-KO hPSC lines. (A) Sequenced genotyping results of *CLDN10*-KO #1 and #2 lines. KO lines #1 and #2 have similar phenotype, both displaying an increased formation of PGC-LC and defective cyst organization. The second coding sequence of *CLDN10* was targeted. (B) Representative sequence traces for each mutations.

## ACKNOWLEDGEMENTS

We thank the Wisconsin National Primate Research Center (WNPRC) Veterinary, Scientific Protocol Implementation, Pathology and Animal Services staff for providing animal care, and assisting in procedures including breeding, pregnancy monitoring, and sample collection. A special thanks to Michele Schotzko, Sara Shaw and Drs. Heather Simmons and Puja Basu for their help in generating the macaque specimens. The contents of this manuscript are solely the responsibility of the authors and do not represent the official views of the NIH. We thank Dr. Deborah Gumucio for insightful comments to the manuscript, as well as the Roy J. Carver Biotechnology Center at the University of Illinois at Urbana-Champaign for sequence services.

## Data and materials availability

All data needed to evaluate the conclusions in the paper are present in the paper and the Supplementary Materials. The raw data, unfiltered count matrix and processed count matrix will be available to the database of Genotypes and Phenotypes (dbGaP) upon publication. Further information and requests for reagents may be directed to Kenichiro Taniguchi.

